# Cross-species epigenomic signatures of exceptional longevity: Insights from Jonathan the Giant Tortoise and human centenarians

**DOI:** 10.64898/2026.11.01.763412

**Authors:** Sara Pischedda, Alberto Gómez-Carballa, Federico Martinón-Torres, Antonio Salas

## Abstract

Can epigenomic signatures of exceptional longevity transcend species boundaries? By comparing findings from a previously published epigenomic study of Jonathan, a 194-year-old giant tortoise, based on buccal scrape and saliva samples, with the analyses of publicly available DNA methylation data from exceptionally long-lived humans based on peripheral blood mononuclear cells (PBMCs), this comparative study reveals intriguing molecular parallels across evolutionarily distant species. Human individuals aged 110+ years exhibited significantly higher global methylation entropy than those aged 60-82 years. Furthermore, 12 candidate genes displayed a distinctive promoter methylation entropy profile, with lower values at the youngest and oldest ages than in intermediate-age groups. Strikingly, these genes were enriched in telomere-related biological processes, highlighting a compelling link between epigenomic patterns and pathways implicated in aging. The correspondence between the findings in humans and previously reported observations in Jonathan suggests testable hypotheses about epigenomic signatures of exceptional longevity that may be conserved across species and extend beyond tissue-specific effects.

---

Very recently, Vaisvil et al. (*1*) reported the first epigenomic characterization of an exceptionally long-lived terrestrial animal, Jonathan, a 194-year-old Aldabra giant tortoise (*Aldabrachelys gigantea*). Their study revealed distinctive DNA methylation patterns associated with extreme longevity, including regions of low methylation entropy enriched in the promoters of genes involved in mitochondrial electron transport and RNA metabolism. These findings provide a unique opportunity to investigate whether epigenomic features associated with exceptional longevity may also be observed across distantly related species. Human lifespan reflects the interplay between intrinsic biological factors and extrinsic influences. Shenhar et al.(*2*) showed that the heritability of human lifespan exceeds 50% when confounding by extrinsic mortality is addressed, highlighting the importance of distinguishing inherited biological determinants from environmental influences. DNA methylation may provide a molecular framework for investigating how environmental exposures and biological processes interact in aging and longevity. Although Vaisvil et al. discussed potential parallels between their findings and previously reported human aging signatures, direct comparisons with human epigenomic datasets are needed to identify shared patterns and generate testable hypotheses about the molecular mechanisms underlying exceptional longevity.

A potentially informative human dataset was reported by Dowrey et al. (*3*), who characterized DNA methylation in peripheral blood mononuclear cells (PBMCs) from a cohort comprising centenarians and their offspring, including individuals described as functionally independent and resilient to disability and cognitive impairment. This dataset provides an opportunity to explore whether methylation features identified in Jonathan are also observed in humans who reach advanced ages.

In the present study, findings from the previously published epigenomic study of Jonathan (*1*), based on buccal scrape and saliva samples, were compared with publicly available DNA methylation data from human PBMCs in the cohort described by Dowrey et al. (GEO accession number: GSE276957). This dataset comprises 25 methylation profiles generated using the Illumina Infinium MethylationEPIC v2 array. The cohort included nine individuals aged 60-82 years, seven aged 100-104 years, seven aged 105-109 years, and two in the 110+ age group (*3*). We first estimated biological age using Horvath’s epigenetic clock (*4*). Estimated biological age was strongly correlated with chronological age across the cohort (Spearman’s ρ = 0.882, *P*-value = 5.8 × 10^−9^, *n* = 25); **Figure 1A**. However, within age groups, the correlation between chronological and estimated biological age was weaker among centenarians (ρ = 0.615, *P*-value = 0.011) than among individuals in the 60-82 age group (ρ = 0.817, *P*-value = 0.007). This attenuation of the linear association at the oldest ages suggests a weaker relationship between chronological age and algorithm-estimated biological age, potentially reflecting a deceleration of estimated biological aging. Nevertheless, this interpretation remains preliminary, as differences in correlation coefficients do not directly establish differences in aging rates.

**Figure 1.**
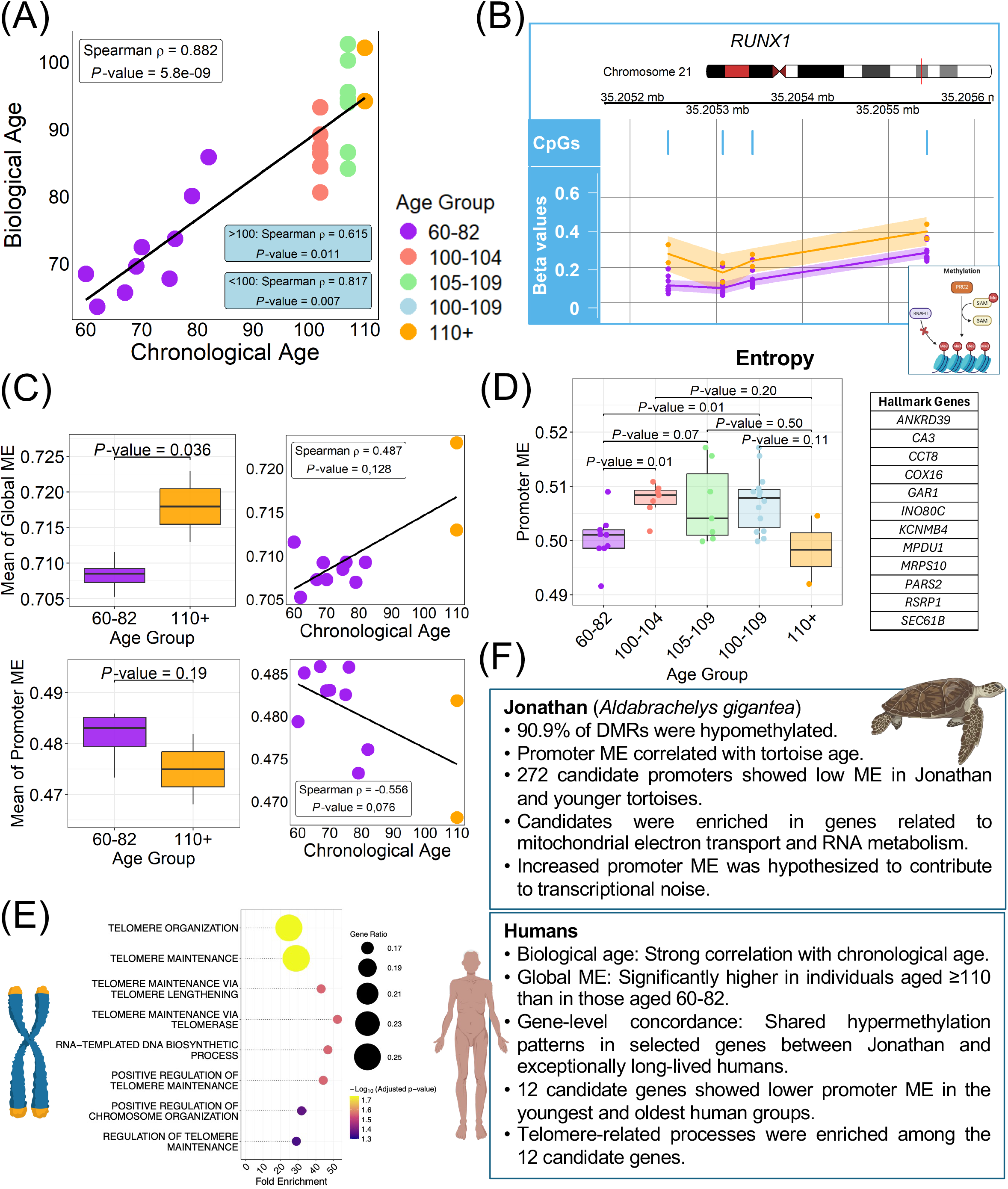
Epigenetic age, DNA methylation entropy, and candidate longevity-associated signatures in humans. **(A)** Relationship between chronological age and biological age estimated using the Horvath clock (*4*). The overall association was strong (Spearman’s ρ = 0.882, *P*-value = 5.8 × 10^−9^). Spearman’s correlation coefficient was lower among centenarians (aged ≥ 100 years; ρ = 0.615) than among the 60-82 age group (ρ= 0.817), suggesting a weaker association between chronological and estimated biological age at advanced ages. (B) Representative example of a hypermethylated DMR within the *RUNX1* gene in centenarians that was also found to be hypermethylated in Jonathan (see **Supplementary Text** for additional examples). Methylation patterns were compared between individuals aged 60-82 years (purple) and those aged 110+ years (orange). Blue vertical marks indicate the genomic positions of CpG sites within the region. Chromosomal locations and genomic coordinates are based on the GRCh38/hg38 reference genome. (C) Global and promoter DNA methylation entropy (ME) in exceptionally long-lived humans. The left panels show the distributions of mean global methylation entropy (top) and promoter methylation entropy (bottom) in the 110+ age group and those aged 60-82 years. The right panels show the relationships between biological age and global methylation entropy (top) and promoter methylation entropy (bottom), with fitted linear trend lines. Orange and purple represent individuals of the 110+ group and the 60-82 age group, respectively. (D) Promoter ME profiles of candidate longevity-associated genes across human age groups. Boxplots show the distribution of individual mean ME values across the 12 selected genes for each age group. (E) Gene Ontology (GO) enrichment analysis of the 12 selected genes, highlighting the overrepresentation of telomere-related biological processes. (F) Summary of the main findings of methylation patterns observed in Jonathan and long-lived humans. A few selected illustrative elements were created using BioRender (https://www.biorender.com).

We identified differentially methylated regions (DMRs) by comparing the 110+ age group with each of the three younger age groups. We then characterized their genomic distribution and direction of methylation change. DMRs were classified as hypermethylated or hypomethylated according to the sign of the mean methylation difference between the 110+ group and the corresponding comparison group.

Hypomethylated DMRs predominated across most genomic categories in the human comparisons, broadly echoing the predominance of hypomethylation reported in Jonathan, where 90.9% of DMRs were hypomethylated (**Supplementary Text**). However, this pattern was less pronounced in humans and varied by genomic region, with hypermethylation predominating in promoters and exons in the comparison between individuals 110+ and the group 105-109 years (**Supplementary Text**). Thus, the human data show partial concordance with Jonathan’s methylation profile, while also revealing differences in how methylation changes are distributed across the genome.

To investigate whether Jonathan’s methylation profile had parallels in human longevity, we examined genes associated with his hypermethylated DMRs in centenarians *vs*. non-centenarians and in the 110+ group *vs*. those aged 60-82 years (**Table S1; Supplementary Text**). Both comparisons revealed hypermethylated DMRs in several genes also identified among those associated with hypermethylated DMRs in Jonathan and annotated as targets of Polycomb repressive complex 2 (*PRC2*), including *LHX8, IRX6, MKX, GDNF*, and members of the NKX family. In the 110+ *vs*. 60-82 years comparison, additional examples included *GATA4, GATA2*, and *RUNX1* (**Figure 1B**). The recurrence of hypermethylation in several genes associated with Jonathan’s profile suggests that some gene-level methylation features may be shared between this exceptionally long-lived tortoise and humans reaching advanced ages.

We also compared methylation entropy between human age groups. Consistent with the age-related pattern reported in Jonathan, global ME was significantly higher in the 110+ group than in individuals aged 60-82 years (*P*-value = 0.036; **Figure 1C**). In contrast, promoter ME showed a lower median in the exceptionally long-lived 110+ group, although this difference did not reach statistical significance (*P*-value = 0.19; **Figure 1C**). Correlation analyses revealed a positive, but non-significant, association between chronological age and global ME (Spearman’s ρ = 0.487, *P*-value = 0.128) and a negative, non-significant association between chronological age and promoter ME (ρ = −0.556, *P*-value = 0.075); **Figure 1C**. Although these correlations should be interpreted cautiously, particularly given the small cohort, their direction is consistent with the contrasting patterns observed for global and promoter ME across age groups (**Supplementary Text**).

Vaisvil et al. identified 272 gene promoters as potential longevity-associated candidates based on low ME in Jonathan and the younger tortoises but higher ME in the two older animals. Building on the analysis performed by Vaisvil et al. we investigated whether the same pattern could help prioritize candidate genes in humans. We examined the corresponding promoter regions in the human cohort and identified 12 of the 272 genes (*ANKRD39, CA3, CCT8, COX16, GAR1, INO80C, KCNMB4, MPDU1, MRPS10, PARS2, RSRP1*, and *SEC61B*) that showed lower mean promoter ME in both the 60-82-year and the 110+ groups than in the intermediate-age groups (100-104,105-109 years, and the combined 100-109-year groups; **Figure 1D**; **Supplementary Text**). The distribution of mean ME across individuals revealed an age-group pattern despite substantial interindividual variability. Promoter ME differed significantly between the 60-82 and 100-104 age groups, as well as between the 60-82 and combined 100-109 age groups (*P*-value = 0.01 for both comparisons). Although the comparison between the 60-82 and 110+ groups did not reach statistical significance, possibly owing to the limited sample size of the 110+ group, the observed distribution suggested a distinct pattern compared with the intermediate-age groups.

These 12 genes therefore represent a focused set of candidates in which the promoter ME pattern originally associated with longevity in Jonathan is also observed in humans at the youngest and oldest ages examined.

Strikingly, Gene Ontology (GO) enrichment analysis of this small set of 12 genes revealed a significant overrepresentation of telomere-related biological processes, including telomere maintenance, telomere organization, telomerase-mediated telomere maintenance, and regulation of telomere maintenance (**Figure 1E**). This result is particularly noteworthy because telomere biology is closely implicated in cellular aging and lifespan regulation (*5*). The emergence of these pathways from a small set of genes selected independently based on their promoter ME patterns provides a compelling lead for further investigation.

The present findings reveal several potential points of concordance between Jonathan and exceptionally long-lived humans, including hypermethylation in selected genes, higher global ME in the oldest human age group, and similar promoter ME patterns in a subset of candidate genes (**Figure 1F**). Notably, these parallels emerge from findings based on different biological sample types: buccal scrapes and saliva in Jonathan, and PBMCs in humans. The observation of concordant patterns across these distinct tissues is particularly intriguing, as it raises the possibility that some epigenomic features associated with exceptional longevity may extend beyond tissue-specific effects. Conversely, differences in tissue composition and cell-type-specific methylation profiles may partly account for the incomplete correspondence between the two species. The enrichment of telomere-related processes adds further biological relevance. Together, these findings raise the possibility that some molecular principles governing aging and longevity transcend species boundaries. Comparative epigenomic studies of organisms with vastly different evolutionary histories and lifespans may help uncover conserved biological mechanisms underlying exceptional longevity, while distinguishing them from species-specific adaptations.

## Consent for publication

All participants have given permission to the publication of the project’s findings.

## Data availability statement

There is no new data generated in the present study.

## Acknowledgements

This study received support by i) Instituto de Salud Carlos III (ISCIII): MANTRA-ID: PI25/00725; PNEUMO-ID360: DTS25/00083; TRINEO: PI22/00162; [to A.S.]; PNEU-MO-V-AI: PI25/01405; OMI-COVI-VAC: PI22/00406; [to F.M.-T.]; NIRSE-omic: PI24/00771 [to S.P.], and co-funded by the European Union; ii) GAIN: IN607B 2020/08 and IN607A 2023/02 [to A.S.]; GEN-COVID: IN845D 2020/23 and IIN607A2021/05 [to F.M.-T.] and IN677D 2024/06 [to A.G.-C.]; iii) ACIS: BI-BACVIR PRIS-3, CovidPhy SA304C, PneumoTrack PRIST-VAL [to A.S.] and Respisal PRIST-VAL [to F.M.-T.]; iv) Consorcio Centro de Investigación Biomédica en Red de Enfermedades Respiratorias (CB21/06/00103) [to F.M.-T. and A.S.]; and v) Spanish Ministry of Science and Innovation (MCIN)/Spanish Research Agency (AEI) and co-funded by the European Union: KAWA-TesT: PID2022-142156OB-I00 [to A.G.-C.]. AGC is supported by the Miguel Servet from ISCIII (CP23/00080) contract, funded by the Instituto de Salud Carlos III (ISCIII) and co-funded by the European Union. The funders were not involved in the study design, collection, analysis, interpretation of data, the writing of this article, or the decision to submit it for publication.

## Authors contribution

AP, AGC, and AS contributed to the conceptualization, methodology, investigation, data analysis, and writing of the manuscript. AS and FMT contributed to funding acquisition. All authors reviewed and approved the final version of the manuscript.

## Supplementary Material

**Supplementary text**. Detailed methodology and results.

**Table S1.** DNA methylation patterns of genes associated with Jonathan’s hypermethylated differentially methylated regions (DMRs) in human longevity comparisons. The table lists genes associated with Jonathan’s hypermethylated DMRs and summarizes the number and direction of DMRs identified in two human comparisons: centenarians *vs*. non-centenarians and the group 110+ *vs*. those aged 60-82 years. For each comparison, hypermethylated and hypomethylated DMR counts, total DMR numbers, and genomic locations are reported. The recurrence of hypermethylated DMRs across multiple genes in the human comparisons highlights similarities with the methylation pattern described in Jonathan.

