## Supplementary Text for "Cross-species epigenomic signatures of exceptional longevity: Insights from Jonathan the Giant Tortoise and human centenarians"

### 1. Detailed Methods

#### DNA methylation preprocessing analysis

DNA methylation data processing and statistical analyses were performed in R using Bioconductor packages. Raw Illumina IDAT files for the GSE276957 dataset were downloaded from the Gene Expression Omnibus (GEO) database using the *GEOquery* package (1). Raw methylation intensity data were imported into R using the *minfi* package (2), and β-values and M-values were calculated. β-values represent the proportion of methylated signal relative to the combined methylated and unmethylated signals and range from 0 to 1. In contrast, M-values represent the $\log_2$ ratio of methylated to unmethylated signal intensities. β-values were used to describe methylation levels and visualize sample distributions, whereas M-values were used for differential DNA methylation. Quality control procedures were applied to assess sample and probe reliability.

Background correction and normalization were performed using the *preprocessQuantile* function from the *minfi* package. This procedure was used to reduce technical variation and address differences in the distributions of methylation measurements across samples and probe designs.

Several filtering steps were applied to obtain a high-quality set of CpG sites. Probes that failed the detection criterion were removed, as were probes located on the X or Y chromosomes and probes affected by known single-nucleotide polymorphisms (SNPs). Finally, cross-reactive probes were excluded using a published list of problematic probes (*3*).

Probe annotations were obtained from the Illumina Human Methylation EPIC v2 annotation package, using the GRCh38/hg38 reference genome.

#### Epigenetic age estimation

DNA methylation-based age estimates were calculated using the Horvath epigenetic clock implemented in the *wateRmelon* package (*4*). The resulting estimates were added to the sample phenotype metadata and used to characterize the study population.

### Identification of differentially methylated regions

Differentially methylated regions (DMRs) were identified using the *DMRcate* package (*5*). To investigate regional DNA methylation differences associated with extreme longevity, three pairwise comparisons were performed: individuals of the 110+ group *vs*. individuals aged 60-82, 100-104, and 105-109 years. These comparisons were designed to characterize methylation differences between the oldest participants and progressively older comparison groups.

DMR analyses were performed using the normalized M-value matrix and a linear model design that included age group and sex.

The identified regions were characterized according to their statistical significance, number of contributing CpG sites, and mean methylation difference (*meandiff*). DMRs were classified as hypermethylated or hypomethylated according to the direction of the mean difference in each contrast. Positive values indicated higher methylation in the 110+ group than in the corresponding comparison group, whereas negative values indicated lower methylation.

Identified DMRs were annotated using the *ChIPseeker* package (*6*) and the *TxDb.Hsapiens.UCSC.hg38.knownGene* transcript annotation database.

Genomic annotations were then grouped into the following broader categories: promoter, exon, intron, untranslated region (UTR), downstream, distal intergenic, and other regions.

### DNA methylation entropy analysis

DNA methylation entropy (ME) was assessed using a Shannon entropy-based approach adapted to methylation β-values, and informed by the conceptual framework described by Chan et al. (*7*).

First, global ME was calculated for each CpG site in each sample using normalized β-values. A promoter-associated ME analysis was performed to investigate entropy specifically at CpG sites annotated as promoter-associated according to the *Regulatory_Feature_Group* annotation of the Illumina manifest file. The same binary Shannon entropy formula was applied to the corresponding β-values, and the entropy values were averaged across the selected CpG sites for each sample.

Associations between ME and chronological age were assessed using correlation analyses. Spearman's rank correlation coefficient was used to evaluate monotonic relationships.

Boxplots were used to visualize the distribution of mean global and promoter-associated ME across the age groups of interest, with particular emphasis on the comparison between individuals aged 60-82 years and the 110+ group. A correlation plot was also generated to evaluate the correlation between ME and chronological age. Statistical significance between groups was assessed using the Wilcoxon test.

### Methylation entropy analysis of candidate genes

The 272 candidate genes listed in Table S8 of the original study by Vaisvil et al. (*8*) were investigated in the human cohort. Gene annotations were integrated with the normalized DNA methylation β-value matrix to identify CpG sites assigned to each candidate gene. Shannon entropy was calculated for each CpG site in each individual, and gene-level mean ME was obtained by averaging the entropy values across the annotated CpG sites for each gene and individual.

Mean ME was calculated separately for the 60-82, 100-104, 105-109, and 110+ age groups. The two intermediate-age groups were also combined into a single 100-109-year category, including all individuals from both groups. Candidate genes were selected according to the following criteria: mean ME in both the 60-82-year and 110+ year groups had to be lower than the corresponding mean ME in each of the 100-104, 105-109, and combined 100-109-year groups. Genes satisfying all these criteria were retained for subsequent analyses. Finally, an individual-level ME score was calculated by averaging gene-level ME values across the selected genes, allowing the distribution of this measure to be examined across age groups.

### 2. Detailed Results

#### Quality control

Quality control of the GSE276957 dataset was performed using the *minfi* package. The methylation intensity profiles were broadly consistent across samples (**Figure S1A**). The beta-value density distributions showed a similar bimodal pattern across samples, with peaks corresponding to low and high methylation levels, respectively (**Figure S1B**), indicating no obvious sample-level deviations in these global methylation profiles.

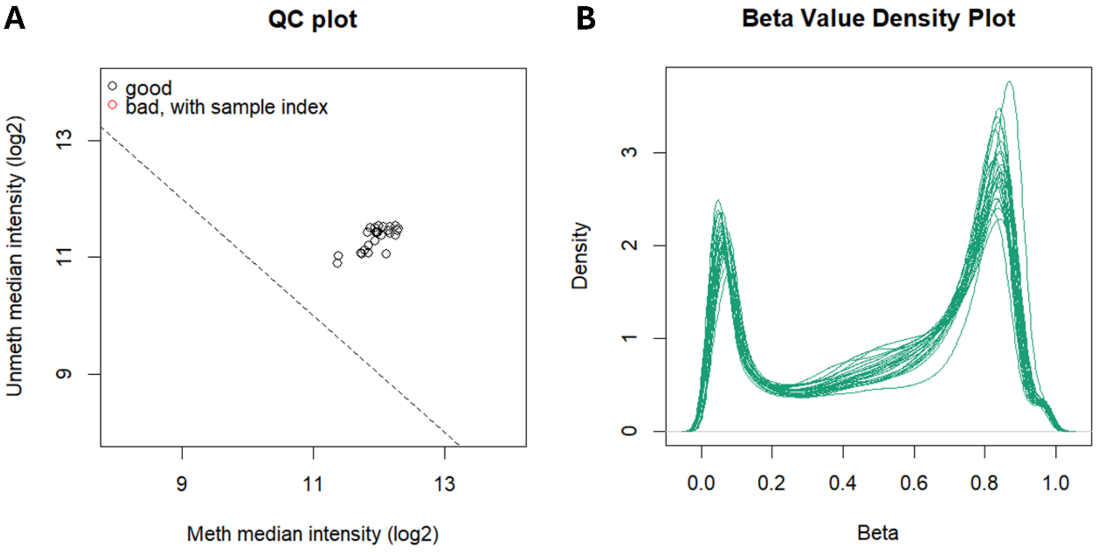


**Figure S1. Quality control of DNA methylation data using the *minfi* package.** (A) Quality control plot showing the relationship between methylated and unmethylated median intensities across samples. (B) Density distributions of DNA methylation beta values across samples, showing a broadly consistent bimodal pattern with peaks at low and high methylation levels.

#### Distribution of differentially methylated regions across genomic annotations in exceptionally long-lived humans

The largest number of significant differentially methylated regions (DMRs) was observed in the comparison between individuals 110+ and those aged 60-82 years. We next examined the distribution of hypermethylated and hypomethylated DMRs across genomic annotations to assess the extent to which the patterns observed in Jonathan were also apparent in humans (**Figure S2**).

Across the three comparisons, hypomethylated DMRs predominated in most genomic categories, although the relative proportions varied across annotations and age-group comparisons. In the comparison between individuals 110+ and those aged 60-82 years, hypomethylated DMRs accounted for 52.4-60.0% of regions across the six annotated categories. The highest proportion was observed in downstream regions (60.0%), followed by untranslated regions (UTRs; 59.5%). In the comparison with individuals aged 100-104 years, hypomethylation predominated in five of the six categories, accounting for 50.8-58.5% of DMRs. Downstream regions were the exception, with hypermethylated DMRs accounting for 69.2%. In the comparison with individuals aged 105-109 years, the distribution was more heterogeneous: hypomethylation predominated in distal intergenic regions (51.3%) and UTRs (56.9%), whereas hypermethylation was more frequent in exons (60.9%) and promoter regions (55.1%). Intronic regions showed a nearly balanced distribution, with 48.8% of DMRs classified as hypomethylated and 51.2% as hypermethylated.

The human data showed a predominance of hypomethylated DMRs across most genomic annotations, broadly consistent with the direction of the changes reported in Jonathan, in which 90.9% of DMRs were hypomethylated relative to the juvenile tortoise. However, the predominance of hypomethylation was considerably less pronounced in humans, and the balance between hypermethylated and hypomethylated DMRs varied across genomic regions. This difference was particularly apparent in promoter and exon regions, where hypermethylated DMRs predominated in the comparison between the 110+ group and those aged 105-109 years. These findings suggest that, although hypomethylation may be a shared feature of the methylation differences observed in exceptionally long-lived individuals across the two species, the genomic distribution and relative prevalence of these changes differ.

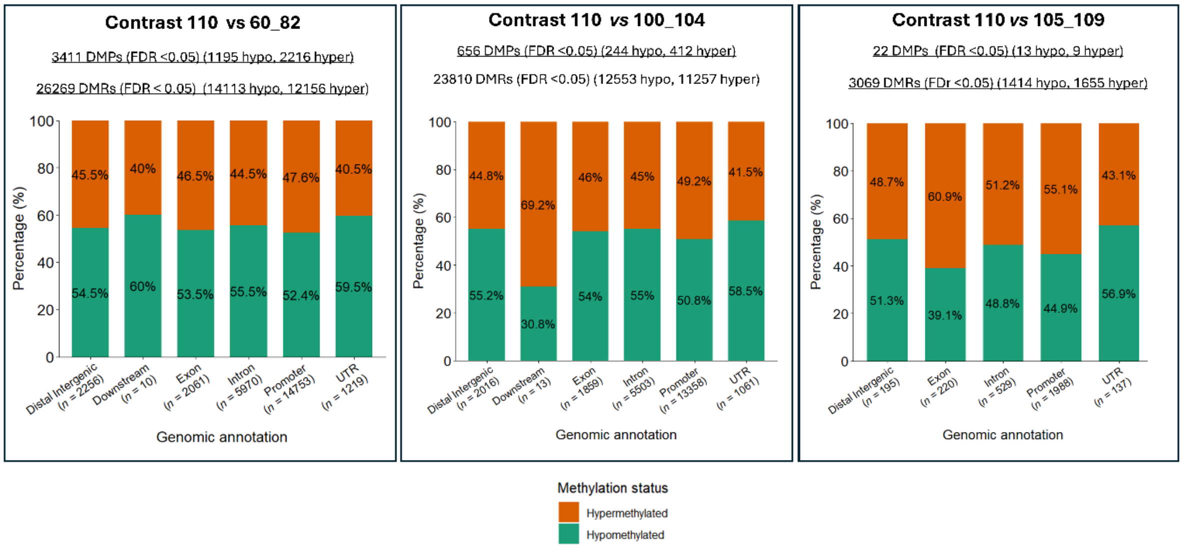


**Figure S2.** Distribution of differentially methylated regions (DMRs) across six genomic annotation categories (distal intergenic, downstream, exon, intron, promoter, and untranslated region [UTR]). The percentages of hypomethylated and hypermethylated DMRs are shown for each of the three contrasts.

### DNA methylation patterns of Jonathan's hypermethylated DMR-associated genes in exceptionally long-lived humans

The analysis of Jonathan's methylome identified 218 genes encompassed by hypermethylated differentially methylated regions (DMRs), including genes involved in developmental pathways and several targets of polycomb repressive complex 2 (PRC2). Among the latter were homeobox genes, including *LHX8*, *IRX6*, *MKX*, and *NKX1-2*, as well as transcription factor genes such as *GATA4*, *RUNX1*, *FOXC2*, *TBX22*, and *TCF21*. To investigate whether these findings had parallels in human longevity, we examined the methylation patterns of the genes listed in Table S7 of the original study in two different comparisons: centenarians *vs*. non-centenarians and the 110+ group *vs*. those aged 60–82 years. We focused on the direction and distribution of DMRs across these groups to assess the extent of concordance with Jonathan's hypermethylation profile (**Table S1**).

In both human comparisons, several genes associated with Jonathan's hypermethylated DMRs also showed hypermethylation, supporting an important degree of concordance across species and longevity groups. In the comparison between the 110+ group and those aged 60-82 years, notable examples included *GATA4*, with five hypermethylated *DMRs*, and *RUNX1*, with seven hypermethylated and three hypomethylated DMRs, respectively. Additional hypermethylation was observed in *GATA2* (three DMRs) and *SHH* (three hypermethylated and one hypomethylated DMR). Other genes, including *LHX8*, *IRX6*, *MKX*, *GDNF*, ISL1, *ISL2*, *NKX2-6*, *NKX2-8*, and *NKX3-2*, also showed hypermethylated DMRs. The centenarian *vs*. non-centenarian comparison likewise revealed hypermethylated DMRs in several genes highlighted in Jonathan's analysis, including *LHX8*, *IRX6*, *MKX*, *GDNF*, *ISL1*, *ISL2*, and members of the *NKX* family. Together, these observations indicate that hypermethylation affecting genes identified in Jonathan is also detectable in both human comparisons (**Figure S3**).

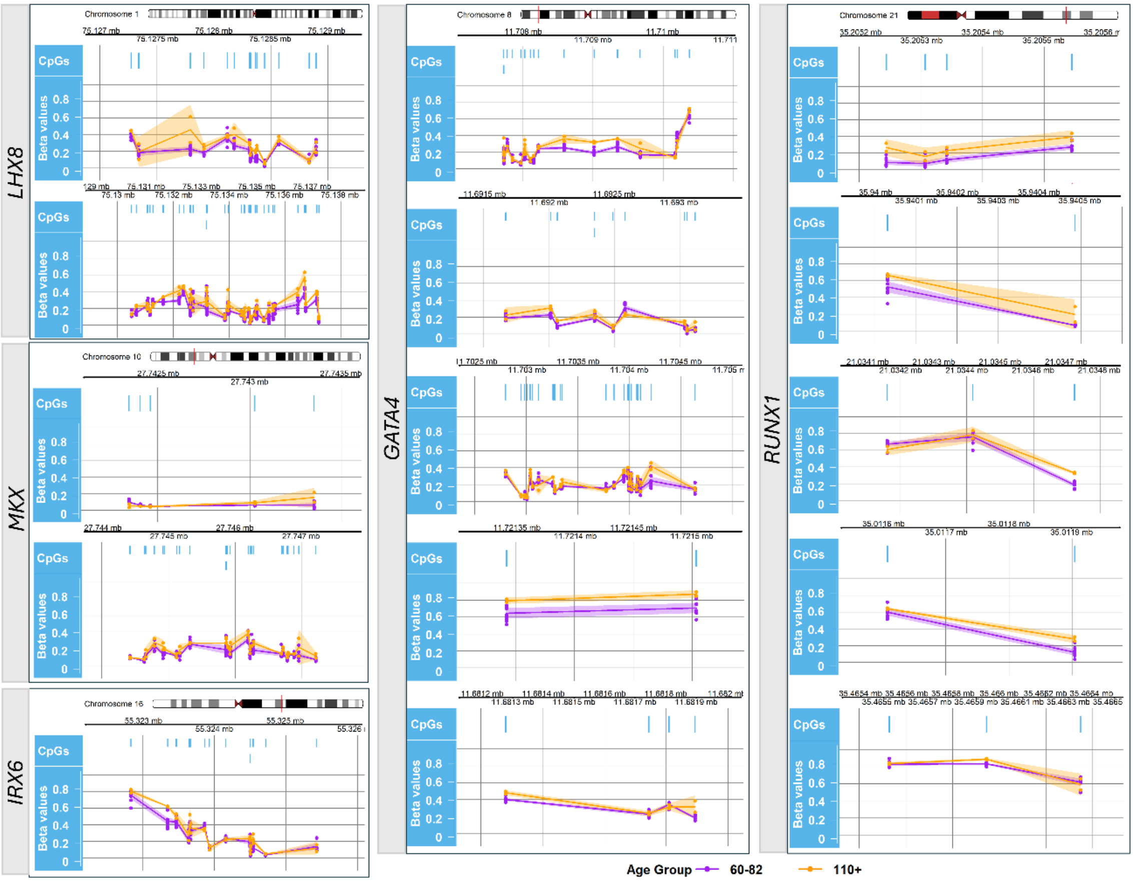


**Figure S3. DNA methylation patterns of selected genes previously associated with hypermethylated DMRs in Jonathan's methylome.** DNA methylation β-values are shown for selected genomic regions of *LHX8, MKX*, *IRX6*, *GATA4*, and *RUNX1*, representing a subset of genes reported in the original study of Jonathan's methylome. Methylation patterns are compared between individuals aged 60-82 years (purple) and 110+ years (orange). Blue vertical marks indicate the genomic positions of CpG sites within each region. Chromosomal locations and genomic coordinates are based on the GRCh38/hg38 reference genome.

The affected regions spanned several genomic features, including promoters, exons, introns, and UTRs. Several genes showed promoter-associated hypermethylation, whereas others exhibited DMRs within gene bodies or UTRs, indicating that the observed similarities were not confined to a single genomic feature. The direction of methylation changes also showed substantial agreement with Jonathan's profile, particularly through the recurrence of hypermethylation across multiple genes and genomic regions.

Some genes exhibited both hypermethylated and hypomethylated DMRs (**Table S1**). For example, in the comparison between the 110+ group and those aged 60-82 years, *WNT5B* and *ZBTB20* each showed two hypermethylated and two hypomethylated DMRs, while *CELSR1* showed one hypermethylated and four hypomethylated DMRs. By contrast, *NKX1-2*, *FZD2*, and *JAG2* showed hypomethylated DMRs without corresponding hypermethylated DMRs in this comparison. Mixed patterns were also observed in the centenarian comparison, including hypomethylated DMRs in *ZBTB20* and *GATA2*. Nevertheless, multiple genes showed hypermethylation in one or both human comparisons, indicating that the overall correspondence was not driven by a single gene. Not all genes reported in the original study had corresponding DMRs in the human analyses.

Therefore, these comparisons suggest similarities between Jonathan's methylation profile and the patterns observed in long-lived humans, both at the gene level and in the direction of methylation changes. The identification of hypermethylated DMRs in multiple genes across the centenarian and exceptionally long-lived comparisons supports the recurrence of this pattern in humans. Although the overlap does not necessarily extend to the same individual genomic regions, the convergence of findings strengthens the evidence for shared DNA methylation features associated with human longevity.

### DNA methylation entropy in exceptionally long-lived humans

DNA methylation entropy in Jonathan, a measure of the disorder or randomness of DNA methylation patterns, was higher than in younger Aldabra giant tortoises across the genome, whereas promoter regions showed a distinct pattern, with low ME across all five tortoises (*8*). Our results in humans reveal a similar age-related pattern, with differences in global and promoter methylation entropy between exceptionally long-lived individuals and younger adults.

Global ME was significantly higher in the 110+ group than in the group of individuals aged 60-82 years (*P*-value = 0.036). This finding is consistent with the higher genome-wide ME reported in Jonathan relative to younger tortoises and suggests that greater methylation disorder may be observed in exceptionally long-lived individuals across species. In contrast, promoter ME was lower in the 110+ years group than in the 60-82-year group, although this difference did not reach statistical significance (*P*-value = 0.19). This distinct promoter pattern is noteworthy in light of the original study, in which promoter regions also differed from the genome-wide pattern and exhibited comparatively low ME across the Aldabra tortoises. Thus, both datasets suggest that the relationship between age and methylation entropy may depend on the genomic region considered.

Correlation analyses were consistent with these observations. Global ME showed a positive association with chronological age (Spearman's $\rho$ = 0.487, *P*-value = 0.128), whereas promoter ME showed a negative association (Spearman's $\rho$ = -0.556, *P*-value = 0.076). Although neither correlation reached statistical significance, the direction of both trends was consistent with the differences observed between age groups: global ME tended to increase with age, while promoter ME tended to decrease. The relatively small sample size may have limited the statistical power to detect these associations.

### Concordance of promoter methylation entropy patterns between Jonathan and exceptionally long-lived humans

In Jonathan, the original study (*8*) identified 272 gene promoters as potential candidates associated with longevity. These promoters exhibited low mean ME in

Jonathan and the younger tortoises (Hatchling and Smalls), but higher ME in the two older animals (Toka and Poka) (*8*). This selection was motivated by the hypothesis that increased promoter ME may contribute to transcriptional noise, whereas the maintenance of low promoter ME in Jonathan could be associated with longevity. To investigate whether these candidate genes might also be relevant to human longevity, we examined the corresponding promoter regions in the human cohort comprising individuals aged 60-82, 100-104, 105-109, the combined 100–109-year groups, and the group 110+.

For each individual, Shannon ME was calculated at individual CpG sites annotated to the 272 genes identified in Jonathan. CpG-level ME values were then averaged for each gene within each individual to obtain a gene-level mean methylation entropy for each gene and individual.

Mean ME was calculated separately for individuals aged 60-82, 100-104, 105-109, and 110+ years. Additionally, the two intermediate-age groups were combined into a single 100-109-year group, including all individuals from both categories. Candidate genes were selected if their mean ME was lower in both the 60-82-year and the 110+ groups than in each intermediate-age group separately and in the combined 100-109-year group.

Twelve genes met these criteria: *ANKRD39*, *CA3*, *CCT8*, *COX16*, *GAR1*, *INO80C*, *KCNMB4*, *MPDU1*, *MRPS10*, *PARS2*, *RSRP1*, and *SEC61B*. These genes represent a subset of the 272 candidates originally identified in Jonathan and display a comparable non-linear age-associated ME pattern in humans, with lower promoter ME in the youngest and exceptionally long-lived groups than in the intermediate-age groups. Given the rationale underlying the original selection in Jonathan, these 12 genes may represent particularly interesting candidates for investigating potentially shared epigenetic features associated with longevity across the two species.

### Limitations of the study

This study has several limitations. First, the human cohort was small, particularly the group 110+. Second, ages for centenarians were provided as intervals (100-104 and 105-109 years) rather than as exact values for each participant. Representative values were therefore assigned to these intervals for correlation analyses, which may have introduced measurement error and affected the estimated strength of associations between age and ME. Additionally, the cross-species comparison involved different biological sample types: buccal scrape and saliva samples in Jonathan, and PBMCs in humans. Although the observed similarities across these distinct tissues are intriguing and may suggest patterns extending beyond tissue-specific effects, tissue composition and cell-type-specific methylation profiles may partly explain the incomplete correspondence between species. Finally, ME was calculated differently in the human and tortoise datasets because the studies used different technologies. In Jonathan, ME was estimated from bisulfite sequencing data using the "me" command in *Metheor*, which captures methylation patterns across sequencing reads, whereas in the human cohort, ME was calculated from array-derived β-values using a site-level Shannon entropy measure. Accordingly, similarities in ME profiles across species should be interpreted as preliminary patterns rather than as evidence that entropy values are directly comparable.
